# A practical pipeline for volume rendering of trillion-voxel tomographic data

**DOI:** 10.64898/2026.04.11.717885

**Authors:** Yusuke Takeda, Daichi Obinata, Takahiro Harada, Mehmet Oguz Derin, Shin Ikegami, Aya Kubota, Shintaro Sasaki, Ryota Fukai, Tomohiro Usui, Kazuki Tainaka, Yasuhiro Iba

## Abstract

Recent advancements in tomography produce imaging data of geological materials (rocks and fossils) at trillion-voxel scales with multi-channels. Such high-resolution datasets are potentially keys to unveil evolutionary biological information with various shapes and sizes that have not been ever discovered. Volume rendering is an ideal visualization approach for them because it treats all voxels without relying on user-defined surface boundaries. However, these large-scale real-world tomographic data have rarely been volume-rendered at their native resolution, limiting the examination of rich morphological information. Here, we demonstrate a de facto standard volume-rendering pipeline running on a graphical processing unit (GPU)-equipped supercomputing system toward multi-channel, trillion-voxel tomographic data. Our workflow preserves original resolution, capturing detailed morphological information spanning microscopic to macroscopic scales. Systematic comparison of node types shows that GPU memory, rather than host memory, is the primary bottleneck. Our results establish a baseline for large-scale, multi-channel volume rendering of real tomographic data and demonstrate its applicability to geological samples. This work is presented as a practical demonstration of large-scale volume visualization.

## Introduction

Tomography is an imaging technique that converts physical specimens into a series of cross-sectional images. It enables the visualization of the internal microstructures, otherwise invisible by the human sense of vision. The major tomographic approaches are non-destructive X-ray computed tomography (CT) and magnetic resonance imaging (MRI), both of which have been utilized in biology to investigate anatomical features [1]. Earth and planetary sciences, especially paleontology and mineralogy, have developed destructive approaches (grinding tomography) with improvements in instruments such as fully automated workflows and higher-resolution image sensors, yielding ever larger datasets with increasing numbers of voxels and color channels in microscopic resolution [2–9]. Recently, digital fossil-mining, a concept to convert whole rock samples into high-fidelity tomographic images and to digitally discover fossils that have previously been overlooked, has been proposed based on the technique [6]. It achieved mass and flawless digital extraction of fragile, morphologically complex fossils which have been difficult to discover and physically extract from hard rocks.

Three-dimensional (3D) visualization of tomography data is an essential process for science research. Broadly, tomographic datasets can be visualized using either surface rendering or volume rendering [10]. Surface rendering displays hollow polygonal 3D models, at the expense of any visual details that lie between extracted surfaces. Volume rendering, by contrast, applies color and transparency to every voxel, retaining the entire dataset. The visualization approach strongly suits destructive tomography that can capture multi-channel images. However, volume rendering remains underutilized for detailed, particularly in large-scale, high-fidelity datasets. It is because the rendering method has mainly focused on grayscale X-ray CT techniques [11,12] and it is only within the last decade that it has become possible to acquire large-scale multi-channel tomographic data using destructive methods [2–8,13,14]. In practical use-cases, volume rendering has been often applied to small-sized data for quick overviews prior to detailed analysis. Large-scale data have been reduced by cropping or resampling, to fit the capabilities of conventional software and lab workstations for volume rendering. The applicability of advanced rendering algorithms for large-scale multi-channel tomographic data remains largely unexplored. Those algorithms, especially in 2010s, were based on the hardware that predates current multi graphical processing unit (GPU) supercomputing environments. Consequently, their reported performance characteristics do not directly reflect the computational regime of today’s large-scale, multichannel tomographic data. The aim of this study is to examine the feasibility and practical constraints of volume rendering for real-world, large-scale, multi-channel tomographic data using a modern supercomputing environment. We achieved volume rendering of trillion-order voxels in full color with red, green, blue, and alpha (RGBA) channels by multi-node parallel rendering and remote visualization. Our case study suggests volume rendering as a powerful technique for discovering hidden structures in large-scale data.

## Background

### Large-scale tomographic images and volume rendering practices

Modern tomographic methods produce volumes exceeding billions to trillions of voxels. For instance, standard X-ray CT can produce 16 billion voxels from 1,000 cross-sectional images each measuring 4000 × 4000 pixels [15]. Synchrotron X-ray CT data can reach ≈ 890 billion (13600 × 13600 × 480) voxels in an extreme case [16]. With the light sheet microscopy, imaging of the full body mouse resulted in as large as 30 billion voxels [17]. High-throughput serial section electron microscopy produces trillion-order grayscale voxels [18]. The method is used to extract specific known structures such as cellular synapses and neurons through image segmentation, so the volume rendering of the whole data has not been presented. Artificially generated grayscale tomographic data also reaches 1 trillion voxels [19,20].

Grayscale tomographic data with moderate number (several billions) of voxels have been processed with both commercial software and freeware, with GPU-equipped workstations. The list of commonly used software for rendering and their limitations can be found in [20,21]. The existing software has a low capacity toward up-to-date large-scale data processing, so the input data is forced to be downsized or trimmed. For example, [22] acquired 15,000 × 14,960 × 5,936 ≈ 1 trillion voxels data with synchrotron radiation laminography (a special case of X-ray CT), but they used downsized 1,875 × 1,870 × 724 voxels data for volume rendering visualization with Dragonfly (Comet Technologies Canada Inc.). Their hardware specification was not described. A volume visualization system for serial sectioning electron microscopic images was proposed and applied it to a trillion (21,494 × 25,790 × 1,850) voxels with three 12-core dual-CPU (3 GHz) machines, 48 GB, with NVIDIA Quadro 6000 (6GB VRAM), but the detailed structure was not visualized sufficiently in 3D because they utilized noisy data [23]. 21,924 × 25,790 × 1,850 voxels grayscale tomographic dataset of mouse cortex acquired by microtome and electron microscopy was visualized in [24], using the same hardware configuration as [23]. [25] proposed software Voreen (University of Münster), specialized in out-of-core processing and visualization of volume data. It is designed for workstation (a midrange consumer PC with an Intel i5-6500, 16 GB RAM, a single NVIDIA GTX 1060 (6GB VRAM) was used in [25]), and does not support distributed computation, so still limited by the capacity of a single machine.

While supercomputers are widely used for large-scale simulations in physics, engineering, and chemistry, they are less commonly used for real tomographic data in biology. Existing volume rendering pipelines for supercomputer outputs often focus on simulation fields such as fluid flow [26] or particle distributions [27]. [19] addressed volume rendering of tomographic data using various supercomputers equipped with CPU nodes; however, the data were generated by artificially upscaling a smaller grayscale dataset consisting of 3.5 million voxels.

### Grinding tomography

Since late 19th century, full-color tomography has been developed in biology by serial sectioning using microtome destructively (see [28] for the history of invention of microtome). The manipulation of thin sections is labor-intensive and time-consuming, making large-scale digital visualization difficult. The volume tends to be thin and not aligned in Z direction (perpendicular direction to photo images), resulting in planar-shaped distorted volume. Meanwhile, researchers in geology and paleontology have developed another destructive approach called serial grinding (grinding tomography). This technique originated from paleontological studies with fossils embedded in rocks [29] (see [2] for the historical review of the method). It involves iterations of sample polishing with a grinding wheel and photographing the exposed surface of the sample. Since then, the method has been elaborated with the help of automation and computerization technologies, enabling the digitization of large volumes of rocks and fossils into digital tomographic images in high-resolution and full-color [2–8]. The photographing can be done with multiple lighting conditions [7,14]. The resolution of the image reaches 240 million pixels [5,6,8,9], collectively reaching a trillion voxels in RGB channels. Unlike those by microtome, the method generates brick-shaping volume which can contain structures with diverse shapes and sizes.

Among commercial software packages, support for multi-channel data by grinding tomography is variable. For example, OsiriX (Pixmeo SARL) and VGStudio (Volume Graphics GmbH)) do not accept multi-channel images or forcibly convert them into grayscale images. In contrast, Amira (Thermo Fisher Scientific Inc.) and ParaView (Kitware Inc.)) can incidentally run volume rendering of RGB images if the data is represented as RGBA. But because the thick, brick-shaped RGB volumes by grinding tomography are novel, practical know-hows about their visualization have not been accumulated well. In addition, the data size of multi-channel images is larger than that of the grayscale images, making the data handling difficult. Previous studies focused on surface rendering [2,6] or visualized only a portion of downsized dataset [3].

## Visualization material

Two datasets of geological samples obtained by grinding tomography were visualized in this study (Table 1). The datasets are public collections at National Museum of Nature and Science (NMNS), Tokyo, Japan. Each 2D image has ≈ 240 million pixels (width: 19,008 pixels, height: 12,672 pixels) with 24-bit RGB TIFF format, resulting in volumes of ≈ 1 trillion voxels (Table 1). The data resolution ranges ≈ submicron to 10 μm per voxel (Table 1). The images contain fossils and biological structures with various shapes and sizes, all of which are the core objectives in paleontology and evolutionary biology. They make good practices in morphological observation through volume rendering.

**Table 1.** Tomographic datasets visualized in this study.

|  | **Dataset A (limestone)** | **Dataset B (silica sinter)** |
| --- | --- | --- |
| **Dataset accession No.** | NMNS_DS00135_wl_01_0001.tif–4059.tif | NMNS_DS00155_wl_01_0001.tif–5001.tif |
| **Dataset size** | 2.7 TB | 3.3 TB |
| **Voxel dimensions** | $19008 \times 12672 \times 4059$ | $19008 \times 12672 \times 5001$ |
| **No. of voxels** | 0.98 trillion | 1.2 trillion |
| **Voxel size** | $3.38\text{ }\mu\text{m} \times 3.38\text{ }\mu\text{m} \times 10\text{ }\mu\text{m}$ | $0.86\text{ }\mu\text{m} \times 0.86\text{ }\mu\text{m} \times 1\text{ }\mu\text{m}$ |

Dataset A represents a limestone block (≈ 60 mm × 60 mm × 40 mm) from the Cretaceous (≈ 90 million years ago) in northern Japan [30, 31]. The detailed locality and geological age are described in [6]. Fossils of both terrestrial and marine organisms lie within a muddy sediment matrix (Fig 1). The visualization aim of this dataset is to demonstrate non-selective 3D extraction of the numerous fossils by making the matrix transparent. The sample was embedded with black-colored resin before grinding. The images were captured under visible white (polychromatic) light (Fig 1A). Additionally, each section was also imaged under ultraviolet (wavelength: 365 nm) light (Fig 1B) which reveals autofluorescent minerals in certain fossils, enhancing their contrast and color definition. The RGB values of the white light image were visualized by volume rendering. The ultraviolet light image was utilized to set the transparency.

**Fig 1.**
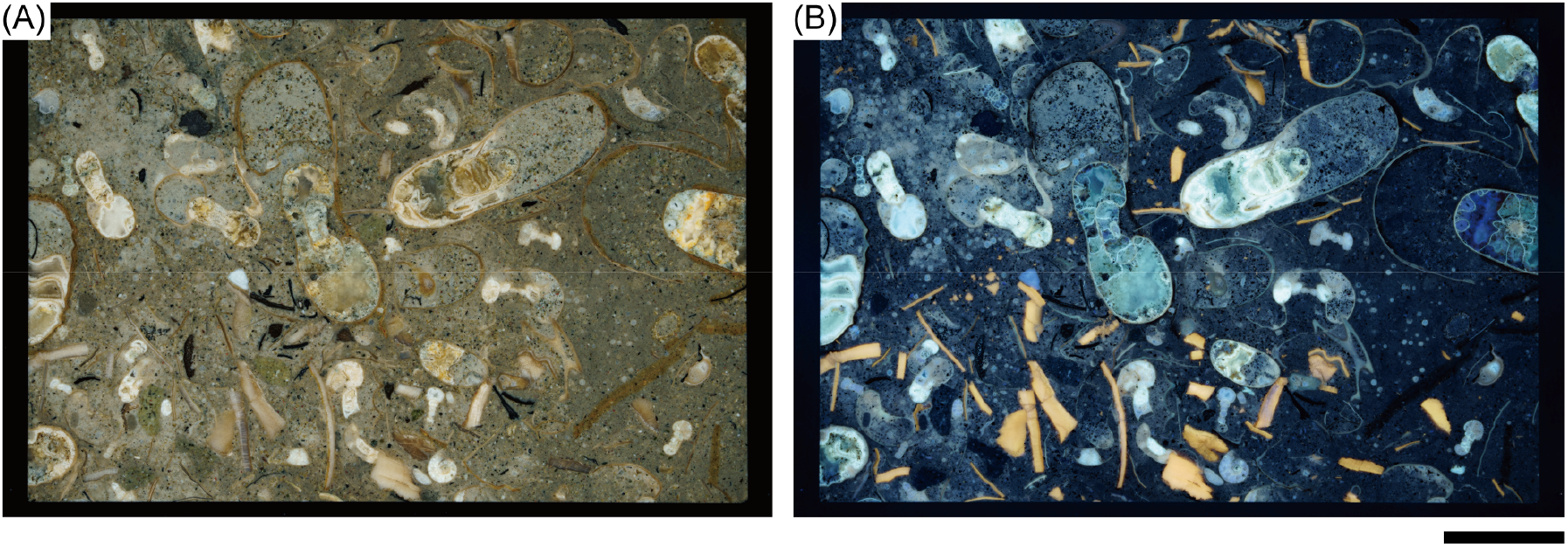
Representative images of dataset A (limestone). (A) An image captured under white light. (B) An additional autofluorescent image captured under ultraviolet light. Scale bar: 10 mm. Original slice images are available from [28].

Dataset B is a silica sinter block (≈ 15 mm × 10 mm × 12 mm) collected from a hot spring in central Japan (Fig 2). The voxel count of the data exceeds 1 trillion (Table 1). It contains fleshly fossilized microorganisms [32], which serve as analogs for early life evolution on Earth. The sample was embedded with black-colored resin before grinding. Then the images were captured under white light.

**Fig 2.**
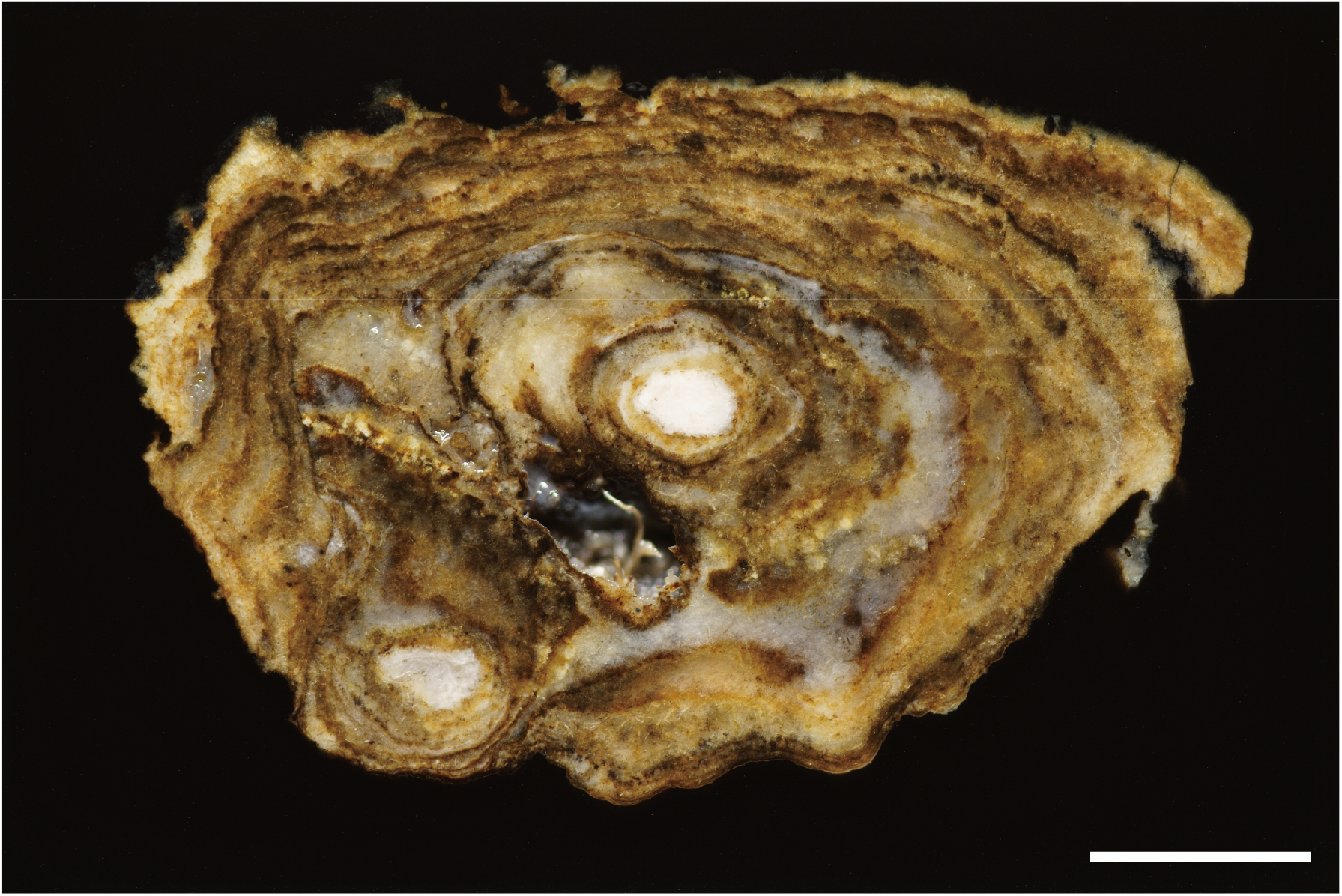
Representative image of dataset B (silica sinter). Scale bar: 10 mm. Original slice image is available from [28].

## Methods

### Visualization method

In this study, we employed a general-purpose visualization software ParaView. We focused on its built-in GPU-based parallel rendering capabilities for large volumes (client server mode; Fig 3, [33]). In this mode, the client runs on a local machine, connecting to a pvserver instance on a supercomputer. Data processing and rendering occur server-side, with user interaction through the graphical interface on the client. Parallel processing of ParaView utilizes Message Passing Interface (MPI), a prevalent techniques in supercomputer systems and computer clusters. After pvserver is invoked through MPI, the data is automatically split into processes. The number of processes can be set independently from the number of nodes. Each process computes volume rendering of each partition using IceT library, a sort-last algorithm for parallel rendering [33,34]. Then the partial rendered images are composited together to form the entire image. Finally, the entire image is displayed on the client machine.

**Fig 3.**
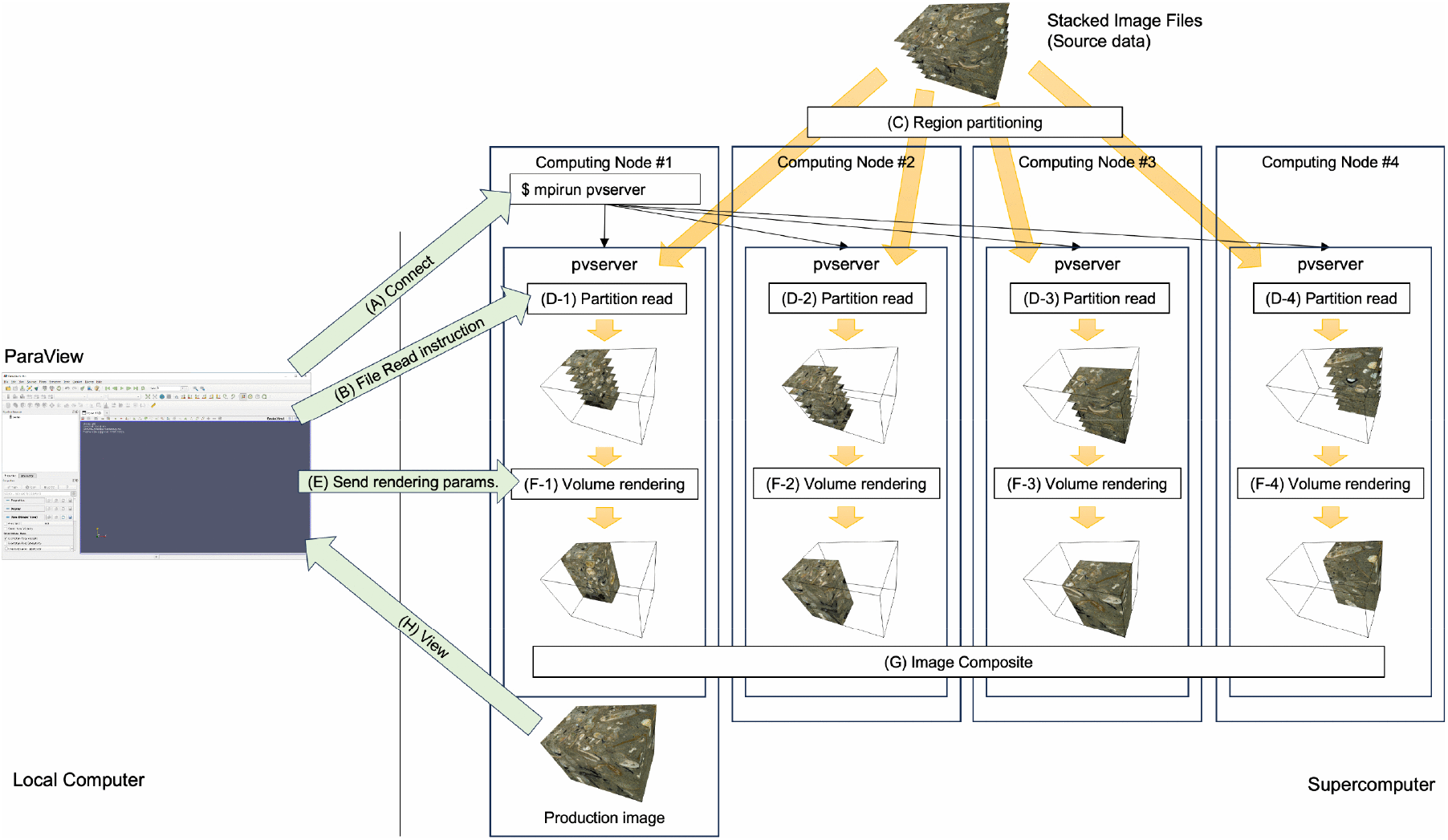
Visualization pipeline for the large-scale volume rendering using ParaView. (A) A MPI parallel job starts and the client machine connected to supercomputer. (B) The supercomputer nodes follow the instruction from the local machine and (C) the input data is partitioned automatically. (D) In each process, the partitioned region image stack is loaded. (E) Following the rendering parameters from the local machine, (F) volume rendering is carried in each process. (G) Finally, the rendered image is composited on the master node and (H) transferred to the client machine.

### Execution environment

All computations in this study ran on the TOKI-RURI system of the Japan Aerospace Exploration Agency (JAXA) Supercomputer System (JSS3). The system is a general-purpose system designed for various computational needs such as visualization and artificial intelligence (AI) [35]. TOKI-RURI is comprised of multiple node types with varying main memory (RAM) capacity and GPU resources per node. We focused on three node types (XM, LM, and ST) sharing the same GPU model but differing in RAM capacity (Table 2).

**Table 2.**
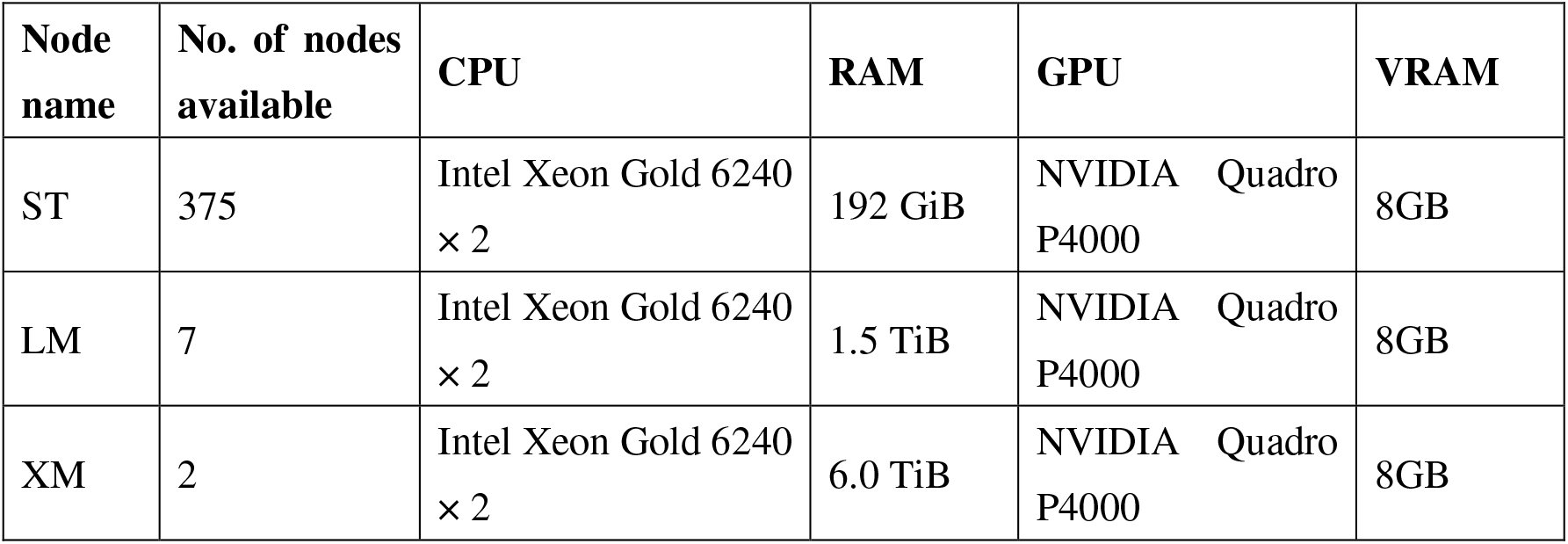
Summary of TOKI-RURI nodes examined in this study.

### Data preprocessing

The TIFF image files were copied to TOKI-FS, the file storage system of TOKI. Since ParaView determines each voxel transparency solely from the alpha channel values baked in the dataset, we generated alpha value from RGB value of each pixel heuristically through a custom Python script, and stored the results in the alpha channel of input images [36]. The computation was primarily based on thresholding. For Dataset A, the surrounding black resin and the sediment matrix were set to fully transparent (alpha = 0), while fossil regions were made partially or fully opaque. Autofluorescent images under UV light facilitated highlighting particular fossil types by subtracting the grayscale value of UV light images from the grayscale value of white light images and applying the result to the alpha channel. For dataset B, the rock region was set to opaque whereas the black-colored surrounding resin was set to transparent.

### Rendering experiments

Volume rendering was executed with ParaView 5.8.1 installed on TOKI-RURI. Since the rendering capability of TOKI-RURI was unknown, we varied the number of nodes and processes per node to identify configurations that successfully handled trillion-voxel volumes. We initially tested smaller volumes (19,008 × 12,672 × 1,000 ≈ 0.24 trillion voxels) using the first 1,000 images of dataset A, and increased the slice count to 2,000, 3,000, and ultimately all 4,048 slices (≈ 0.98 trillion voxels). We examined combinations of node type (XM, LM, or ST), the number of nodes (2–100), and the number of processes per node (1–18). We recorded whether rendering completed successfully and evaluated the visual output for completeness.

## Results

Table 3 summarizes the experimental configurations in this study. Rendering with 2 XM nodes was not carried out correctly even for the smallest dataset (1,000 images), although the node has sufficient main memory size and all 1,000 RGBA images were loaded. The rendering showed only fragmented partial blocks of the volume (Fig 4). Adjusting processes per node did not help. Since there are only 2 nodes available for XM, further tests using XM was not conducted. Similarly, LM nodes could load 1,000 slices when spread across four nodes but failed at 2,000 slices, even when using all seven available nodes.

**Table 3.** Data size, settings for the node use, and the result of volume rendering.

| Voxel dimensions | Node | No. of nodes | No. of process per node | Result |
| --- | --- | --- | --- | --- |
| $19,008 \times 12,672 \times 1,000$ | XM | 2 | 1 | Failed |
| $19,008 \times 12,672 \times 1,000$ | XM | 2 | 18 | Failed |
| $19,008 \times 12,672 \times 1,000$ | LM | 4 | 9 | Succeeded |
| $19,008 \times 12,672 \times 1,000$ | ST | 36 | 1 | Succeeded |
| $19,008 \times 12,672 \times 1,000$ | ST | 50 | 1 | Succeeded |
| $19,008 \times 12,672 \times 2,000$ | LM | 36 | 9 | Failed |
| $19,008 \times 12,672 \times 2,000$ | ST | 36 | 1 | Failed |
| $19,008 \times 12,672 \times 2,000$ | ST | 50 | 1 | Succeeded |
| $19,008 \times 12,672 \times 2,000$ | ST | 100 | 1 | Succeeded |
| $19,008 \times 12,672 \times 3,000$ | ST | 50 | 1 | Failed |
| $19,008 \times 12,672 \times 3,000$ | ST | 100 | 1 | Succeeded |
| $19,008 \times 12,672 \times 4,048$ | ST | 100 | 1 | Succeeded |
| $19,008 \times 12,672 \times 5,001$ | ST | 100 | 1 | Succeeded |

**Fig 4.**
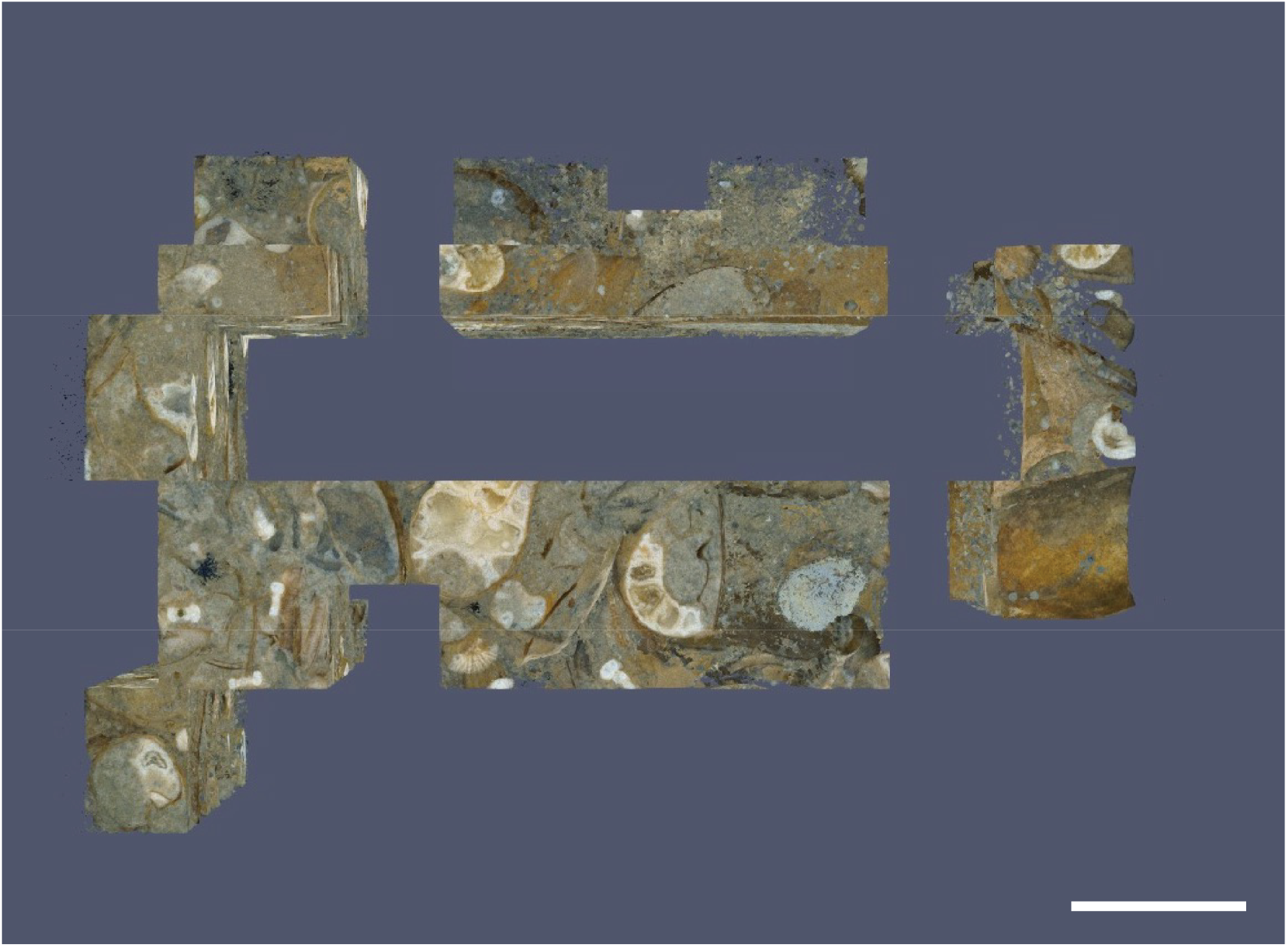
Failure case of volume rendering of dataset A, with 2 XM nodes and 18 processes per node. Scale bar: 10 mm.

In contrast, ST nodes, which have the smallest main memory, handled the larger volumes successfully, provided we allocated enough nodes. Rendering 1,000 slices required 36 ST nodes, while 2,000 slices needed 50, and beyond 3,000 slices demanded 100 nodes (Table 3). With 100 ST nodes and one process per node, we successfully volume-rendered all 4,048 slices of dataset A and saved the output as a 2,128 × 1,666 PNG image (Fig 5). Muddy sediments were effectively made transparent, and fossils were reconstructed in 3D with full-color (Figs 5A–C). Utilization of multi-light sources worked to extract various colored fossils from bright-colored shells (Figs 5A, C, and D) to dark-colored plants (Figs 5B, E). Fecal pellets and bacterial structures that were indistinct in white-light images were also visualized (Figs 5C, F). These structures span in wide range, from micrometer to centimeter scales (Fig 5F).

**Fig 5.**
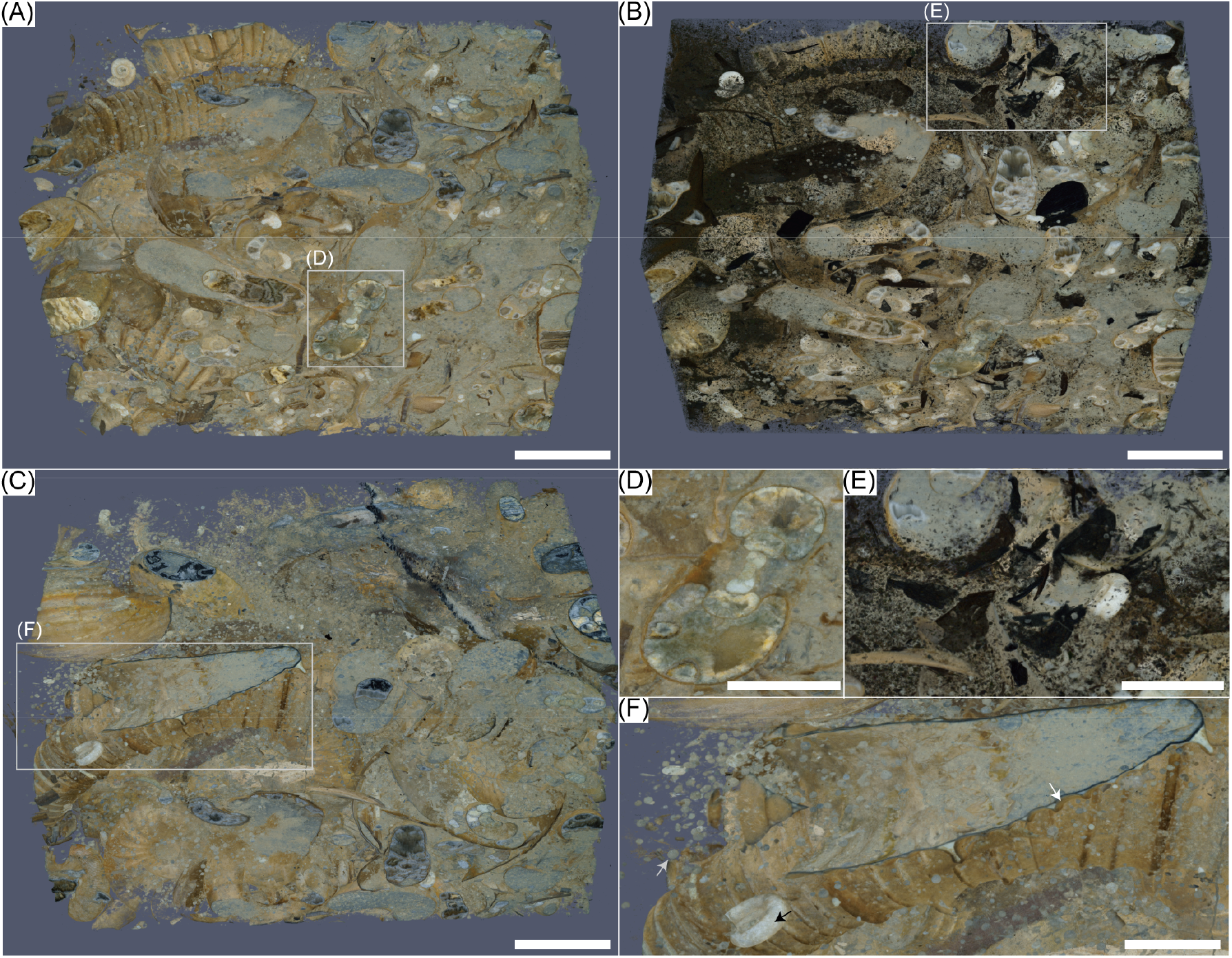
Visualization of dataset A (limestone) using 100 ST nodes on which single process is executed, showing numerous fossils preserved in a volume of 40 mm × 40 mm × 60 mm. (A) Volume rendering of bright-colored fossils. (B) Volume rendering of both bright- and dark-colored fossils. (C) Visualization of the back side of (A). (D) A section of an ammonite fossil with complex internal structure (septa). (E) Fragments of fossil plants (dark-colored materials) which have been difficult to visualize. (F) Various size fossils are preserved in the sample, represented by a centimeter-sized ammonite shell (white arrow), a millimeter-sized juvenile ammonite (black arrow), and micrometer-scale spherical structures probably produced by biological activities such as excretion (gray arrow). Scale bars: 10 mm for (A)–(C) and 5 mm for (D)–(F), respectively. The high-resolution version of the figure is available from [37].

Finally, the volume rendering of dataset B (1.2 trillion voxels) was carried out successfully with the same settings. The morphology of the irregular-shaped rock sample was reconstructed in full-color (Fig 6A). Geobiological structures including layers of mineral deposits and microbial filaments are visualized (Fig 6B).

**Fig 6.**
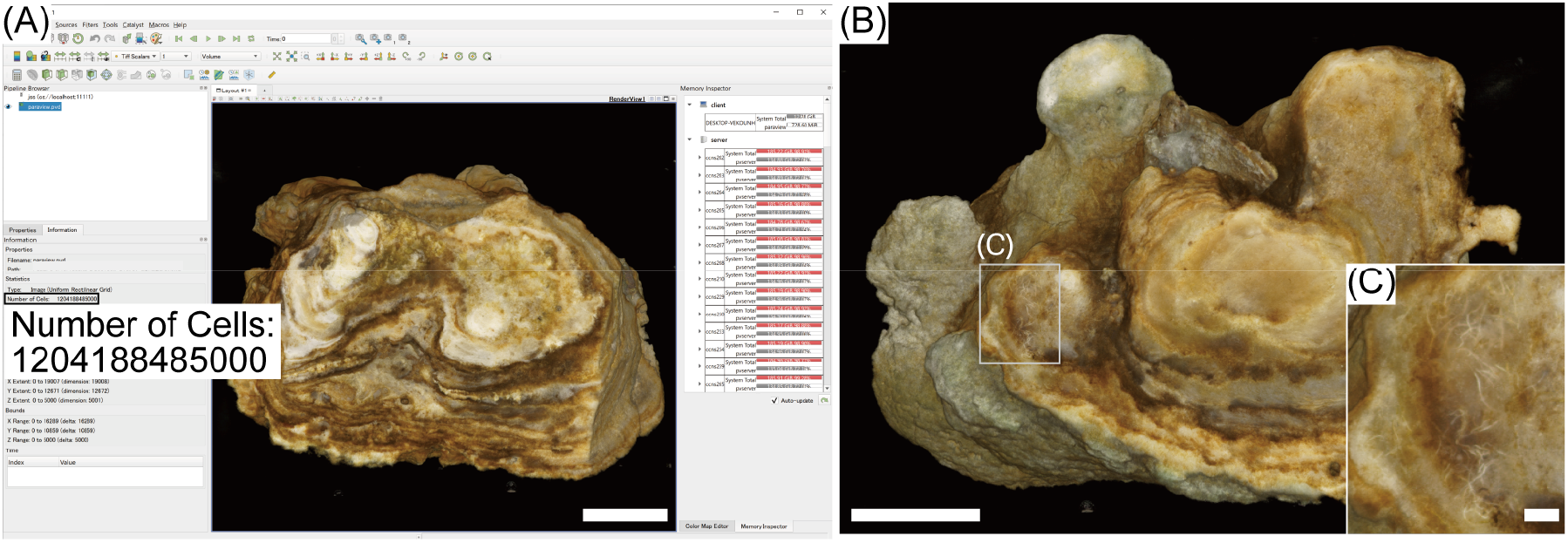
Visualization of dataset B (silica sinter) using 100 ST nodes on which single process is executed. (A) A screenshot of ParaView’s GUI running volume rendering running on local machine, showing the number of cells (voxels) processed exceeds 1 trillion. (B) Visualization of the back side of (A). (C) A close-up view of microbial filament structures. Scale bars: 10 mm for (A) and (B), and 1 mm for (C), respectively. The high-resolution version of the figure is available from [37].

## Discussion

### Large-scale volume rendering of tomographic images on a supercomputing platform

Supercomputers have become user-friendly, offering remote GUIs and streamlined job submission systems. They are becoming more accessible to a diverse range of users. By employing a de facto standard volume rendering pipeline implemented in ParaView, our study demonstrates that supercomputers are effective platforms for visualizing large-scale bioimaging data. Researchers who lack extensive skills in algorithm implementation or powerful local hardware can still render trillion-voxel multi-channel volumes by relying on established software and algorithms.

This study provides practical insights into parallel GPU-based visualization. Many supercomputing systems offer nodes with varying specifications for main memory, CPUs, and GPUs. As in JSS3, two general strategies are possible: using a small number of high-spec nodes (XM), or deploying a large number of low-spec nodes (ST). Our experiments on XM, LM and ST nodes represent such comparison with different main memory size. XM nodes, which have the largest main memory in JSS3-TOKI RURI, failed in volume rendering. Although the main memory of the instance is large enough to store volume data, these nodes could not store the entire data on GPU memory (VRAM). There are only two XM nodes available in the system, not enough for the data size and resulting in incomplete rendering. In contrast, ST nodes, which have the smallest main memory, successfully achieved one-trillion-voxel volume rendering without errors by aggregating GPU memory across tens to hundreds of nodes. Thus, distributing data across a sufficient number of GPUs, even on nodes with relatively small main memory, proved more critical than using a small number of high-memory nodes. These results indicate that the primary bottleneck for large-volume visualization is GPU memory, not CPU performance or host memory capacity, underscoring the importance of multi-GPU scaling when designing future visualization systems and algorithms.

Despite its capability to handle large-scale volume data, the approach in this study has several limitations. Interactivity can be constrained by remote-rendering latency and by the substantial time required to load, process, and composite massive volumes. The large data size also leads to prolonged transfer times from local machines to supercomputer storage. Achieving higher-resolution screenshots would require additional VRAM and, consequently, more GPU nodes. As data sizes increase further, these issues become even more critical. Moreover, for multi-channel (RGB) tomographic data, opacity (alpha channel) assignment currently depends on preprocessing rather than fully interactive, real-time transfer-function manipulation, limiting dynamic control of transparency. Our baseline results therefore highlight where future software and algorithmic developments should be particularly focused on reducing GPU memory pressure, improving data streaming and compositing for remote visualization, and enabling more flexible, interactive handling of multi-channel volume data.

### Revealing hidden fossils through volume rendering

We successfully applied full-color volume rendering to trillion-voxel, multi-channel real tomographic data at their original size using ParaView and a supercomputer. Although our demonstration features fossil-bearing rock samples through grinding tomography, the voxel count and the number of the channels reflect the state-of-the-art in bioimaging, indicating that the approach extends to other large-scale tomographic datasets. Imaging methods such as fluorescence imaging with light sheet microscopy [38], which are originally grayscale but increasingly multi-channel, can also benefit from this approach.

Fossils embedded in hard rock are the only direct physical evidence of deep-time life evolution. They have contributed to the elucidation of early life evolution, diversification, and mass extinction. Precise phylogenetic analysis still requires fossil evidence to time-scaling and accuracy in collaboration with molecular genome information. Tomography-based paleontological studies have focused on the internal structures of target fossils either already exposed on or fully separated from rocks. It has been common practice to label and segment each target fossil specimen prior to three-dimensional visualization. Machine-learning approaches that exploit the morphological characteristics of the target fossils have also been attempted [2]. Those workflows can be applied to rock samples in which the presence of fossils is evident from the sample surface and in which the types of fossils present inside the rock are predictable. However, there also exist rocks, such as dataset A in this study, that contain a heterogeneous assemblage of diverse fossils. In such case, before undertaking individual morphometric analyses, it is necessary to obtain an overview of all fossils present and to clarify what kinds of fossils occur and how they are distributed. Observing cross-sections of fossils in serial two-dimensional images are not enough for humans to recognize fossils that possess complex three-dimensional morphologies. In this study, we successfully visualized almost all fossils embedded within in a fist-sized (≈ 10 cm-scale) rock sample rocks retaining their colors, as shown in Fig 5. Transparency was controlled by thresholding rather than by explicit segmentation of individual fossils. The sizes, types, distributions and orientations of the visualized fossils are highly variable and cannot be predicted from the outer surface of the rock. They provide information on interactions between organisms and their environments, as well as inter-individual interactions (Figs 5 and 6). These results emphasize the value of preserving original data resolution during visualization without downsampling. Our approach also enhances the effectiveness of scientific visualization of non-biological samples with various sized hierarchical structures (Fig 7; [4]).

**Fig 7.**
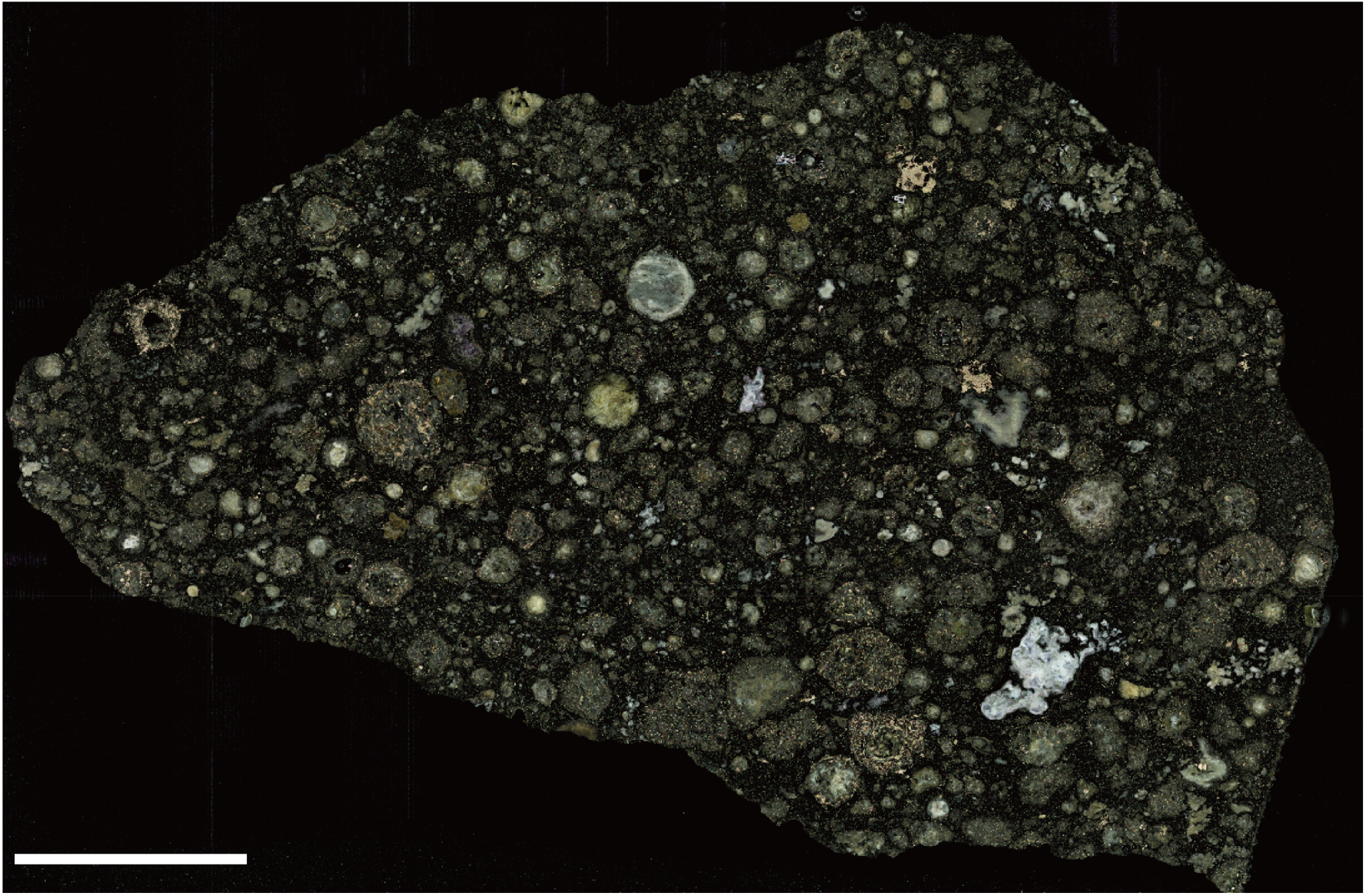
Large-scale volume rendering of non-paleontological materials. Visualization of tomographic images of a meteorite section (Allende, CV3 carbonaceous chondrite; 53685 × 83474 × 20 ≈ 0.09 trillion voxels) with ParaView and JSS3, showing numerous mineralogical inclusions with diverse color, size, shape and texture. The image dataset is public collection at NMNS (accession No. NMNS_DS00162_wl_01_0001.tif–NMNS_DS00162_wl_21_0020.tif). Scale bar: 10 mm.

## Conclusion

This study focused on large-scale volume rendering of tomography data and examined the effectiveness of real-world large-scale visualization in the context of paleontology. We used trillion-voxel, multichannel grinding-tomography data of fossil-bearing rocks, whose data dimensions are comparable to state-of-the-art tomographic techniques. By selecting GPU nodes with sufficient aggregate memory and employing a de facto standard volume-rendering pipeline, we achieved distributed volume rendering of these datasets without downsampling, thereby establishing a practical baseline for large-scale tomographic visualization. Under these conditions, almost all fossils, ranging from microbial to macroscopic organisms, were visualized in situ and in three dimensions, as preserved within the rock matrix. Owing to the nature of volume rendering, both internal and external biological structures were seamlessly extracted at high resolution. These visualizations extend the scope of morphological observation across multiple spatial scales. Our findings underscore the importance of high-fidelity tomographic acquisition and large-scale volume rendering as an effective approach in paleontology, evolutionary biology and related fields.

## Acknowledgments

The use of JSS3 was carried out as JSS3 large-scale challenge program. We thank Takeshi Hoshino, Naoyuki Fujita, Akira Fujioka, and Susumu Takatsu (all in JAXA) for their supports with JSS3. Shintaro Sasaki and Yumi Adachi (all in Hokkaido University) are thanked for imaging experiments. We also thank Tetsuya Yokoyama (Institute of Science Tokyo) for providing the Allende meteorite sample.

## Author contributions statement

Y.T.: conceptualization, investigation, formal analysis, data visualization, funding acquisition, writing—original draft; D.O.: methodology, writing—review and editing; T.H.: validation, writing—review and editing; M.O.D.: validation, writing—review and editing; S.I.: resources, funding acquisition, writing—review and editing; A.K.: curation, writing—review and editing; R.F.: resources, writing—review and editing; T.U.: resources, writing—review and editing; K.T.: resources, funding acquisition, writing—review and editing; Y.I.: project administration, funding acquisition, writing—review and editing. All authors read and approved the final version of the manuscript.

## Data Availability Statement

The tomography image data underlying this article are the collections at National Museum of Nature and Science (NMNS), Tokyo, Japan. The total size of datasets utilized in this study is 6.3 TB. Image examples (2.7 GB) for the validation of this study are available at Figshare (https://doi.org/10.6084/m9.figshare.30931406.)

